# Revisiting the cancer microbiome using PRISM

**DOI:** 10.1101/2025.01.21.634087

**Authors:** Bassel C. Ghaddar, Martin J. Blaser, Subhajyoti De

## Abstract

Recent controversy around the cancer microbiome highlights the need for improved microbial analysis methods for human genomics data. We developed PRISM, a computational approach for precise microorganism identification and decontamination from low-biomass sequencing data. PRISM removes spurious signals and achieves excellent performance when benchmarked on a curated dataset of 62,006 known true- and false-positive taxa. We then use PRISM to detect microbes in 8 cancer types from the CPTAC and TCGA datasets. We identify rich microbiomes in gastrointestinal tract tumors in CPTAC and identify bacteria in a subset of pancreatic tumors that are associated with altered glycoproteomes, more extensive smoking histories, and higher tumor recurrence risk. We find relatively sparse microbes in other cancer types and in TCGA, which we demonstrate may reflect differing sequencing parameters. Overall, PRISM does not replace gold-standard controls, but it enables higher-confidence analyses and reveals tumor-associated microorganisms with potential molecular and clinical significance.

## Introduction

We now know that the human microbiome can impact numerous organ systems^1^. Particular microbiome compositions have been associated with broad aspects of human life, including cardiovascular disease^2^, cancer^3^, autoimmune disease^4^, depression^5^, longevity^6^, fertility^7^, and responses to treatment modalities^8^. Despite increasing research into the human microbiome, however, a number of basic questions remain unanswered, including questions about its size, distribution^9^, and presence in various human tissues, exemplified by recent controversies around the fetal microbiome^10,11^ and cancer microbiome^12,13^.

In the cancer microbiome field, analyses of the same pan-cancer genomic data by independent researchers have reached differing conclusions, ranging from widely abundant to absent microbiomes^12,14–18^. In breast cancer, detection of bacterial lipopolysaccharide (LPS) based on tissue staining^19^ has not been well-reproduced^20^. In pancreatic cancer, 16S ribosomal gene sequencing was found to discriminate short and long-term survivors in one study^21^, but was minimally detected in a separate cohort^22^. Others have observed that some of the cancer-type specific microbes detected in one analysis of The Cancer Genome Atlas (TCGA)^17^ have only been associated with extreme environments and not with humans^12^ and that many reported microbial signals can be attributed to microbial database contamination by human and vector sequences^14^.

The basis of these discordant conclusions can be attributed to both falsely negative and falsely positive results. Falsely negative findings arise from variation in source tissues or insufficient sensitivity for detection; falsely positive results arise from both contamination and technical artifacts that lead to inadequate specificity of putative detections and possible dilution of true signal. Contamination originates from whole microorganisms or adventitious microbial biomatter within reagents or from the ambient environment^23–25^. Technical artifacts arise from incorrect reference genomes, wherein contamination by human, vector, or microbial DNA is a known issue^26,27^, as well as from errors in taxonomic classifiers, which often are only benchmarked using simulated datasets due to a lack of challenging, real-life ground-truth datasets^28,29^. Some of these challenges can be mitigated by using gold-standard contamination controls at each experimental step and using full-length read alignment against high-confidence microbial reference databases. However, contamination controls are often not present, and unbiased, full-length alignment of large datasets against large references (e.g. with BLAST^30^) is computationally infeasible.

As such, there is a clear need for improved analysis methods to identify true (tissue-resident) microbial signals, especially in the low microbiome biomass setting profiled by genomic sequencing. Here we develop PRISM (**Pr**ecise **I**dentification of **S**pecies of the **M**icrobiome), a computationally efficient method to identify and mitigate sources of technical error and contamination in genomic sequencing data. We benchmark PRISM using data from 150 separate studies with known uni-microbial and poly-microbial burdens, and we show that it identifies tissue-resident taxa with high sensitivity and specificity and produces qualitatively appropriate results. We then use PRISM to assess the microbiome in 8 cancer types in the Clinical Proteomic Tumor Analysis Consortium (CPTAC) and in TCGA. We identify subsets of tumors with rich microbial content, find associations of microbial populations with tumor molecular and clinical characteristics, and explore causes of microbial differences between CPTAC and TCGA. PRISM confirms and substantially improves upon our previous work on SAHMI^31^ and pancreatic cancer^32^ with increased rigor, direct interrogation of microbial reads, and large-scale, quantitative benchmarking.

## Results

### Overview of PRISM

The purpose of PRISM is to identify taxa that are truly-present in the source sample and that are not falsely positive artifacts or contaminants. False-positive artifacts are taxa reported by taxonomic classifiers but which are incorrectly classified; the reads assigned to these taxa did not originate from the reported taxa. Contaminant taxa are correctly identified, but these reads were not present in the source tissue; they were introduced during sample handling. PRISM thus has two key conceptual steps: (1) identifying correctly classified taxa and eliminating false positive artifacts, and then (2) distinguishing between truly-present vs. contaminating taxa.

To do this in a computationally efficient manner, PRISM leverages the speed of k-mer based taxonomic classification and the accuracy of full-length alignment. PRISM removes important technical artifacts and identifies uniquely identifiable taxa using full-length mapping of a representative subsample of sequencing reads and taxa first identified using the k-mer-based taxonomic classification. These taxa are correctly classified, but they may have originated from the original tissue (truly-present) or could be contaminants. PRISM then employs a machine learning classifier to predict tissue-present microbes vs. contaminants based on multiple features engineered from read mapping statistics (**Fig. 1a**). As described below, this classifier was trained on a large, curated dataset with known truly-present as well as contaminating taxa, and its predictions can be used to prioritize species in a sample that are highly likely to be validated as truly-present.

**Figure 1.**
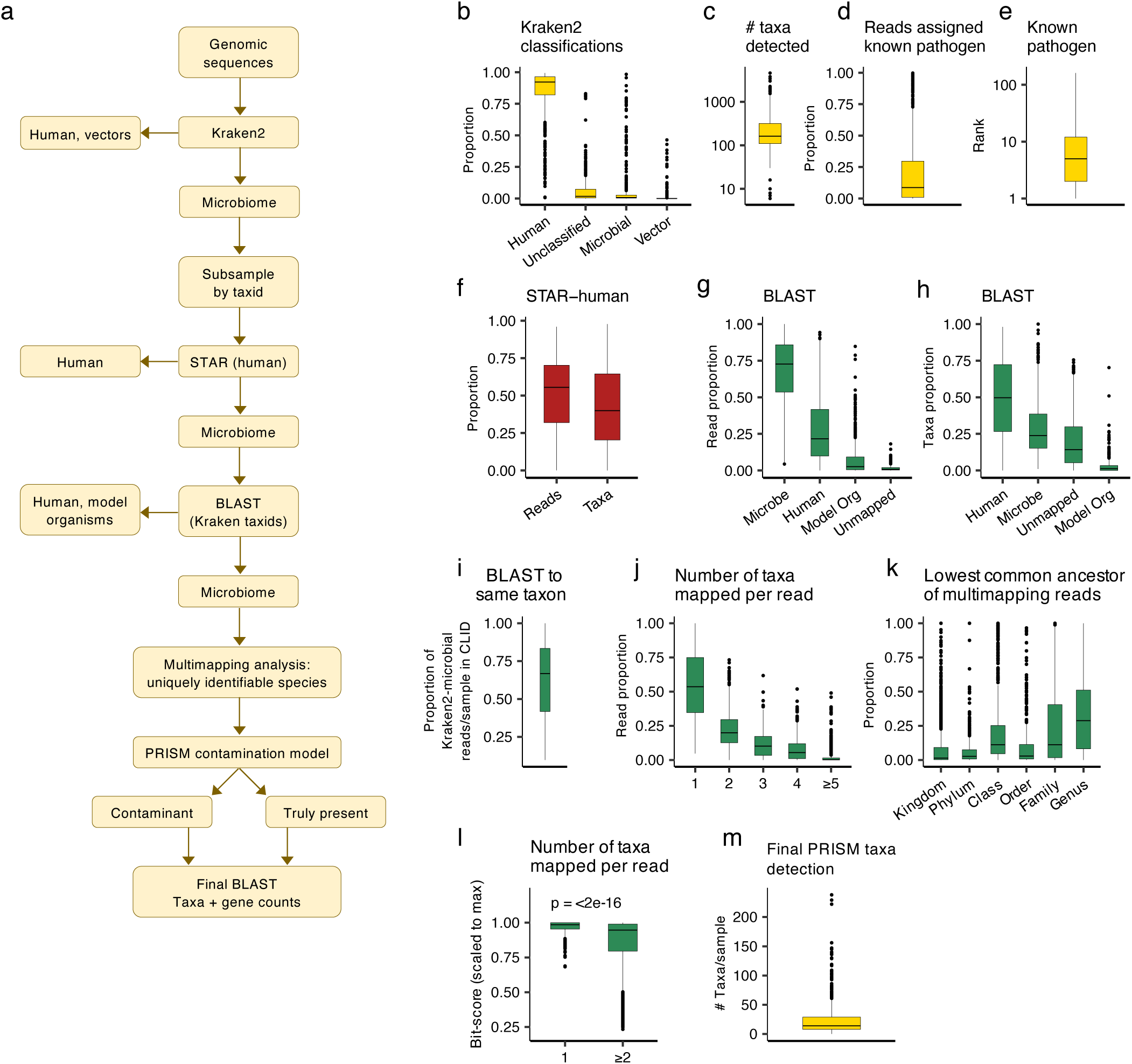
PRISM algorithm for detecting uniquely identifiable taxa. **(a)** Workflow diagram of the key steps in the PRISM algorithm. **(b)** Boxplots of Kraken2 summary results on the cell line infection dataset (CLID; n=677 samples). In all panels, boxplots show median (line), 25^th^ and 75^th^ percentiles (box) and 1.5xIQR (whiskers). Individual points represent outliers. (b) shows proportion of reads assigned to each domain. **(c)** Number of species with >10 reads detected per sample. **(d)** Proportion of species-level reads assigned to the known pathogen in each sample. **(e)** Rank of the known pathogen in sample by read counts, with “1” representing the most abundant. **(f)** Boxplots indicating the proportion of reads classified as microbial by Kraken2 and number of taxa/sample in CLID that were mappable to the human genome by STAR. **(g)** Boxplots indicating the proportion of reads and taxa **(h)** per sample in CLID that mapped to each domain using BLAST. These are reads and taxa that were classified as microbial by Kraken2 and then not Human by STAR. **(i)** Kraken2 + BLAST: Boxplot indicating the proportion of reads per sample in CLID that were mapped to the same microbial taxon by both Kraken2 and BLAST. **(j)** Boxplots indicating the proportion of reads per sample in CLID that mapped to 1 or more taxa with BLAST. X-axis indicates the number of taxa mapped by a given read **(k)** Boxplots indicating proportion of reads per sample in CLID that map to multiple species with BLAST that are resolvable to each lowest common ancestor. **(l)** Boxplots comparing the BLAST bitscore/sample (scaled to the max bitscore in the sample) in CLID between single-mapping and multimapping reads; Wilcoxon testing. **(m)** Boxplots indicating the number of taxa detected by PRISM/sample in the CLID.

For PRISM’s initial development, we compiled a unique dataset of 677 human cell-line RNA-seq experiments from 136 independent Sequence Read Archive (SRA) projects, hereon referred to as the cell line infection dataset (CLID, **Table S1**). These samples had known uni- or poly-microbial infections from 70 unique bacteria, fungi, and viruses. In each sample, the known, live, growing microbe(s) is defined as the microbe that is truly tissue-present, and all other detected taxa are presumed to be contaminants or artifacts. Based on this classification, we next describe the impact of PRISM’s key algorithmic steps toward identifying truly-present microbes in unknown samples.

### Identifying high confidence microbial taxa vs. false positive artifacts

PRISM begins with k-mer-based taxonomic classification using Kraken2^33^ and a reference database of complete microbial (bacterial, fungal, viral) genomes, the human genome, and known vector sequences from RefSeq^34^, as is standard with Kraken2^33^. Across all of the surveyed data from 667 studies in the CLID, Kraken2 classified 87% of reads as human, 5.8% unclassified, 5.4% microbial, and 1.1% as vectors (**Fig. 1b**). It identified the known pathogen in all samples, but it also detected a median of 160 other microbial species per sample (**Fig. 1c**). The known pathogen was assigned only a median of 9% (mean=21%) of the reads classified at the microbial species level (**Fig. 1d**) and on average was ranked the 13^th^ (median=5^th^) most abundant microbial species per sample (**Fig. 1e**). Since these data were from controlled cell-line experiments, the abundance of other taxa was surprising and suggested that a large number of the reads and taxa were misclassified.

To efficiently probe the reads, PRISM next creates a representative subset of the data by randomly sampling reads from each species identified by Kraken2. It then does full-length sequence alignment of the subsampled data, starting with mapping to the human genome using STAR^35^. Despite including the human genome in Kraken2, a median of 55% of Kraken2-identified microbial reads/sample (corresponding to 40% of taxa) in CLID were mapped to the human genome by STAR (**Fig. 1f**). Those reads were removed and the remaining data were then simultaneously mapped to genomes of humans, common model organisms, and the remaining Kraken-identified microbial taxa using BLAST^30^. Per sample, a median of 73% of the putative microbial reads (23% of taxa) aligned to microbial genomes, while 22% of reads (50% of taxa) mapped to the human genome, 2.6% of reads (1.3% of taxa) to model organisms, and 0.8% of reads (14% of taxa) were unmapped (**Fig. 1g-h**). Examining the reads that mapped to human, 25% mapped to ribosomal sequences, 7% to mitochondrial, 4% to CpG island sequences, 3% to chimeric sequences, 2% to HLA genes, and 2% to nuclear genes (**Table S2**). These data clearly identify substantial mapping errors that account for large percentages of putative microbial reads and taxa, many due to missed human reads.

PRISM next removes sequencing reads that map to the human genome and then analyzes the remaining microbial reads. The majority (73%) of Kraken2-identified microbial reads per sample also mapped to a microbial taxon with BLAST; however, only 68% (median) of Kraken2-identified microbial reads mapped to the same taxon in BLAST (**Fig. 1i**). There also was substantial multimapping of reads across microbial taxa; 46% (median) of reads per sample were mappable to >1 taxon (**Fig. 1j**), with 29% (median) of the multimapping reads only resolvable to the “family” taxonomic level or higher (**Fig. 1k**). These multimapping reads had significantly lower mapping scores (BLAST bit-score, relative to sample maximum) compared to reads mapping to only one taxon (Wilcoxon p<2e-16, **Fig. 1l**), indicating they were more repetitive or conserved sequences that were more likely to be mapped by chance.

Because a large number of taxa were attributable only to multi-mapping reads, PRISM next identifies the species that are supported with uniquely mapping reads. Although this approach is conservative, it identified all 870 species that were truly-present in CLID, even those with as few as 10 reads, along with a median of 14 taxa per sample (**Fig. 1m**), a substantial reduction from an initially detected median of 160 taxa/sample (**Fig. 1c**). The other species detected were presumed contaminants, the most common being the human cutaneous species *Cutibacterium acnes* and *Malassezia restricta,* as well as *Escherichia coli, Klebsiella pneumoniae,* and *Staphylococcus aureus* (**Table S3**). Once the uniquely identifiable species are determined, PRISM identifies all reads that can be attributed to them and does a second BLAST to these species to determine final species and gene counts. Although Kraken2 consistently ranks amongst the best taxonomic classifiers^28,29^, this analysis confirm that (1) the majority of the taxa reported from the studies analyzed were false positives due to human and microbial multimapping artifacts and (2) that PRISM reliably detected truly-present taxa.

### Data-driven prediction of truly-present vs. contaminant taxa

After identifying the taxa that are robustly supported by the data, PRISM next uses a machine learning model to distinguish between taxa that are truly-present in tissues and contaminants. To train this model, we assembled a large dataset with four main data types: (1) CLID (n=677 samples, as described above), (2) *in silico* combinations of randomly selected reads from CLID samples to create n=153 new samples each with 4-45 truly-present species and 1-200 contaminants (CLID-C), (3) *in silico* combinations of cultured and whole genome sequenced (WGS) bacterial isolates from human gut^36^, bladder^37^, and the FDA ARGOS database^38^ as well as WGS from negative control samples^39–44^ (n=374 samples; WGS), and (4) n=41 samples from metatranscriptomic data from human gut^45^, mouth^46^, and vagina^47^ (META). In total, our combined dataset contained 1,245 samples, each with 1-133 true positive taxa detected and 10^1^-10^7^ microbial reads detected. PRISM reported 37,052 total observed taxa which consisted of 552 unique true positive species and 2,279 unique contaminant species in this training data, comprising a total of 14,875 taxa that were true-positives and 22,177 that were contaminants. This dataset is unique in its size, *a priori* determination of truly-present, growing microbes vs. contaminants, and composition of multiple datatypes, independent projects, and targeted and nontargeted sequencing approaches. Sample details are in **Table S3**.

We used these data to engineer 5 classes of features (total of 40 variables, see **Methods**) to be used by the PRISM model. The underlying hypothesis behind these features was that sequencing approaches will capture and map a greater diversity of sequencing reads from truly-present species compared to contaminants, and that this diversity could be quantified in multiple complementary ways. The first class of features quantified taxon gene-product diversity using the BLAST results, where we observed that truly-present, growing microbes showed a significantly larger number and diversity of genes and protein products compared to contaminants (Wilcoxon p<2e-16, **Fig. 2a, Fig. S1a**). The second class of features related to the multi-mappability of some reads, where we quantified 3 metrics based on the observation that truly-present taxa had a significantly lower proportion of reads mappable to multiple taxa (Wilcoxon p<2e-16, **Fig. 2b, Fig. S1b**). The third class of features was based on Kraken2’s read count estimates and the observation that truly-present taxa had significantly higher unique read counts (Wilcoxon p<2e-16, **Fig. 2c, Fig. S1c**). The fourth class of features was “k-mer phylogeny”, or the proportion of k-mers per taxon assigned to each taxonomic level by Kraken2, where we observed that truly-present species had significantly more k-mers classified at higher resolutions (order to species) levels (Wilcoxon p<2e-16, **Fig. 2d, Fig. S1d**). The fifth class of features was based on Kraken2 misclassifications, where we observed that when a microbe was truly-present, Kraken2 identified a greater number of phylogenetically related taxa and assigned significantly more reads to them (Wilcoxon p<2e-16, **Fig. 2e, Fig. S1e**). We confirmed that these trends remained significant even while comparing the same taxa when they were truly-present vs. when they were contaminants (**Fig. S1f**) and while controlling for read abundance as well as the number of phylogenetically related taxa present in the reference database (**Fig. S1g**).

**Figure 2.**
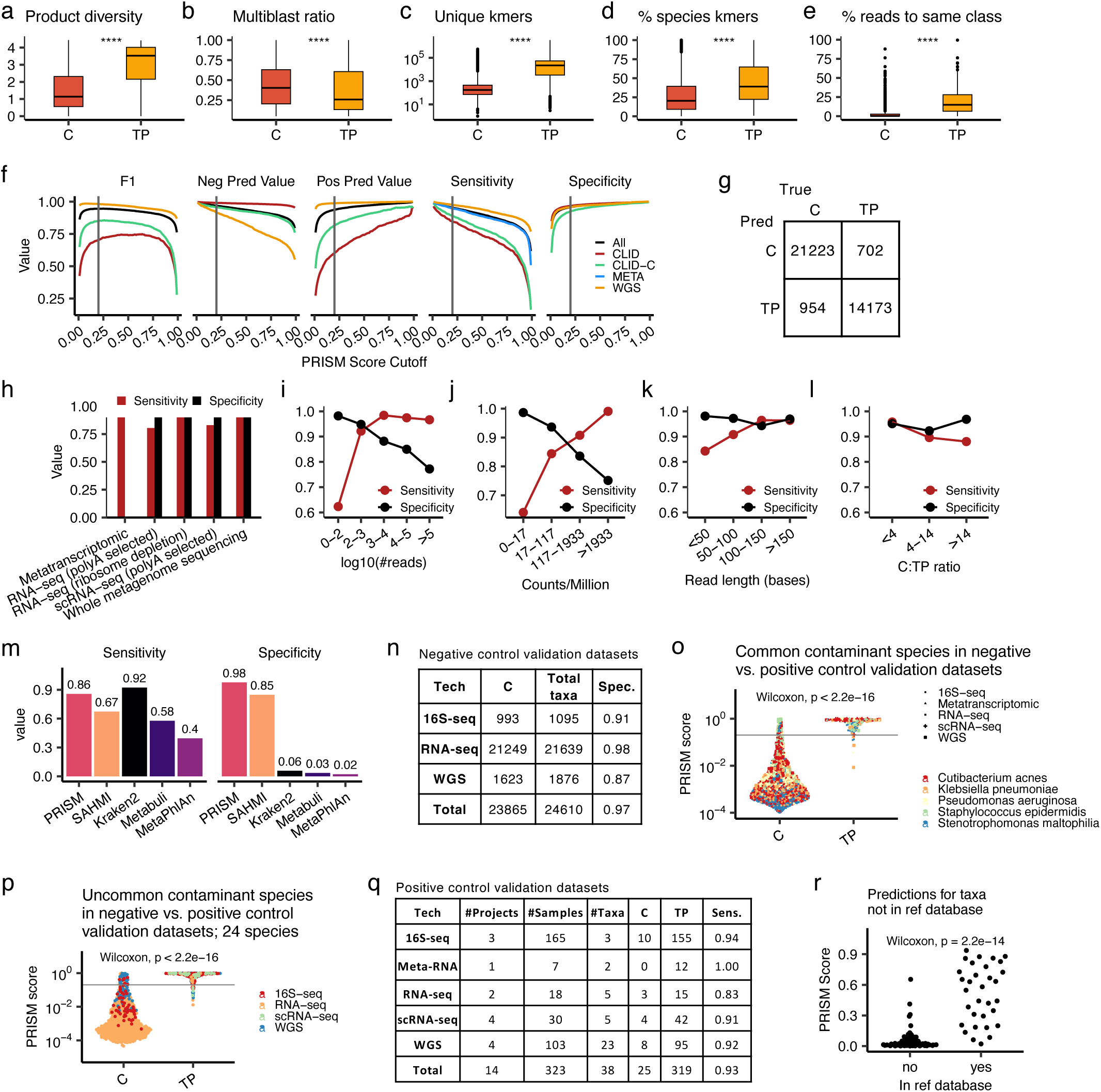
Machine learning model predicts reliable PRISM scores. **(a-f)** Boxplots comparing the value of key PRISM model features between known contaminant (C) taxa and truly-present (TP) taxa in the CLID datasets. See Figure S1 for all features and see Methods for specific feature details and formulas. In all panels, boxplots show median (line), 25^th^ and 75^th^ percentiles (box) and 1.5xIQR (whiskers); points represent outliers. Wilcoxon testing; ****, p<0.0001. **(a)** Microbial product Shannon diversity/taxon. **(b)** Multimapping vs uniquely mapping reads proportion/taxon. **(c)** Unique k-mer counts/taxon. **(d)** Proportion of k-mers/taxon resolvable to the species level. **(e)** Proportion of reads/sample assigned to the same class as the given taxon. **(f)** Performance metrics of PRISM in classifying truly-present vs. contaminant taxa across a range of PRISM score cutoffs. The vertical gray line marks a PRISM score of 0.2, the threshold used throughout the paper. (**g**) Confusion matrix showing the PRISM model performance for independent test datasets in classifying contaminants (C) and truly-present (TP) taxa in the combined CLID, CLID-C, META, and WGS datasets. Data were stratified 5-fold by project of origin for training and testing, i.e. all samples from the same study were assigned to the same fold. **(h-l)** PRISM performance across a range of metrics. **(h)** Bar-plot indicating the PRISM model performance stratified by sequencing type. **(i)** Sensitivity and specificity of the model for taxa with varying numbers of reads. **(j)** Sensitivity and specificity across a range of taxon relative abundance (counts per million total reads). (**k**) Sensitivity and specificity across different sequencing read lengths. **(l)** Sensitivity and specificity across samples with varying ratios of contaminants to truly-present taxa (C:TP). (**m**) Bar plots comparing the sensitivity and specificity of five methods for identifying truly-present taxa in CLID. For methods other than PRISM, genera from a “contaminant blacklist” were additionally filtered from the outputs. (**n**) Table showing PRISM specificity in predicting contaminants from negative control datasets, using PRISM score cutoff of 0.2. Key: C, contaminant; Total taxa, total number of taxa detected; Spec., specificity. (**o**) Swarm plot comparing PRISM scores for five common contaminant species, examining their scores when they were known to be contaminants in the negative control datasets (C) vs. when they were known to be truly-present (TP) in human infections. The grey line marks PRISM score = 0.2. Points are colored by species and shaped by the sequencing type. (**p**) Swarm plot similar to (o) but for 24 species that were not common contaminants in the negative control datasets. (**q**) Table showing PRISM sensitivity across all of the positive control datasets. C, predicted contaminant; TP, predicted to be truly-present; Sens, sensitivity. (**r**) Swarm plot of PRISM scores for samples from the CDC-HAI project that contained WGS of taxa that were (yes, n=20) or were not (no, n=28) in our reference database.

We used our curated datasets and computed features to train a gradient-boosted tree model to classify species that were truly-present or were contaminants. This model predicts a “PRISM score” for each taxon, i.e. the probability that a taxon is a contaminant (0) or is truly-present (1). The model was tested using 5-fold cross-validation, during which information leak was prevented by stratifying the data (including the synthesized samples) by project of origin, ensuring that testing was done on samples from SRA projects unseen by the model during training. The model had excellent performance in classifying truly-present taxa in the independent test projects across a range of data types and PRISM score cutoffs (**Fig. 2f**). To benchmark its performance, we chose a PRISM score cutoff of 0.2, as this marked an elbow in the performance plots where there were marginal gains in the model’s F1 score and specificity at greater values while maintaining higher sensitivity (**Fig. 2f**). Using this cutoff, PRISM had an overall sensitivity=0.95, specificity=0.96, positive prediction value=0.94, negative predictive value=0.97 (McNemar test p<2e-16, **Fig. 2g**). Performance remained high across all sequencing types (**Fig. 2h**; a specificity calculation is not provided for the META (metatranscriptomic) datasets because they only included truely-positive taxa – see Methods), a range of reads per taxon (especially for taxa with >100 reads, **Fig. 2i**), a range of relative abundance (counts per million total reads, **Fig. 2j**), a range of read lengths (especially >100 bases, **Fig. 2k**), and a range of ratios of contaminants to truly-present species (**Fig. 2l**). Lastly, we compared PRISM’s performance on CLID (its lowest performing dataset) to three other high sensitivity or specificity methods for taxonomic classification (Kraken2^33^, MetaPhlAn^48^, Metabuli^49^) and one method for computational decontamination (SAHMI^31^), and we included the additional step of filtering a “blacklist” of 497 common contaminating genera^17^ from their outputs. PRISM was both more sensitive and specific in calling truly-present microbes than all four methods (**Fig. 2m**). These tests demonstrated the reliability of the PRISM score across a range of conditions and its improvement over existing methods.

### Validating PRISM

We next validated the PRISM score using several new datasets and scenarios. First, we tested PRISM on negative control data from RNA-seq, WGS, and 16S rRNA gene sequencing (16S-seq) experiments. For the RNA-seq data, we used cell line samples with no known infection, presuming that all detected microbes are contaminants, and for the WGS and 16S-seq data we used negative control samples (reagent or blank samples); sample details are in **Table S4**. Using the same PRISM score cutoff of 0.2 as above, PRISM had excellent specificity in identifying contaminants across all data types (RNA-seq: 21249/21639 (98%), 16S-seq: 993/1095 (91%); WGS: 1623/1876 (87%); **Fig. 2n**).

Second, using these negative control data, we tested whether the PRISM score had a bias towards taxa that were more frequently present as true-positives in the training data. We binned taxa into deciles by their frequency as true-positives in the training data and examined their PRISM scores in the negative control data. Although there was a statistically significant difference in PRISM scores across the groups (Kruskal-Wallis, p<2e-16, **Fig. S2a**), the median PRISM score across all groups was <0.004 and the 80^th^ percentile was <0.06, indicating that across all of the test groups the majority of taxa were correctly classified without ambiguity. To examine whether the statistical bias was due to increased training instances or to other factors intrinsic to these specific data, we ran three experiments in which we subsampled the training data to include only 10, 15, or 25 instances of each contaminant and truly-present taxon, retrained the model, and predicted new PRISM scores for the negative control validation data. Keeping taxa in the same decile groups based on their frequency in the complete training data (the same groups as in **Fig S2a**), we continued to observe significant differences in PRISM scores in all experiments (Kruskal-Wallis, p<2e-16, **Fig. S2b**), indicating that the differences in PRISM scores across training instance frequency groups were being driven by sample and taxa-specific factors and not by bias related to training data taxon frequency.

Third, we assessed PRISM’s ability to identify species that are truly-present in the specimens but are typically common contaminants. We studied five of the most common contaminants (*Cutibacterium acnes* [681 samples], *Klebsiella pneumoniae* [463 samples], *Pseudomonas aeruginosa* [367 samples], *Staphylococcus epidermidis* [201 samples], *Stenotrophomonas maltophilia* [292 samples]) identified in the negative control datasets. Then, we used PRISM to analyze 7 other datasets^50–56^ in which these species were validated as being truly-present in positive cultures from human infections. PRISM had an overall sensitivity of 0.95 and specificity of 0.97 (from the negative control datasets) in identifying these species (**Fig. S2c**), and their PRISM scores were significantly higher when they were truly-present than when they were contaminants in the negative control datasets (Wilcoxon p<2.2e-16, **Fig. 2o**). This trend was significant for each taxon individually (**Fig. S2d**) and across all tested sequencing types (**Fig. S2e**). The PRISM score also was a more useful classifier than taxon read count. Using a logistic regression model to predict contaminant vs. truly-present as a function of the PRISM score and read counts, the PRISM score p-value was p<2e-16, whereas the read count p-value was p=0.21 (**Fig. S2f**), indicating that the PRISM score added significant value towards identifying truly-present species.

Fourth, we assessed PRISM’s ability to correctly classify species that are uncommon contaminants and to distinguish them when they are truly-present. We identified 24 species in the negative control validation datasets that were present in fewer than 10% of samples and we then analyzed 6 additional sequencing studies in which these species were validated as truly-present based on positive cultures from human infections. These positive control datasets included infections of human stomach with *Helicobacter pylori*^57^, skin with *Mycobacterium leprae*^54^, lung with SARS-CoV-2^58^, colon cancers colonized by *Fusobacterium nucleatum*^59,60^, and several organisms causing bacteremia^61^, urinary tract infections^62^, or infections in other internal body fluids^56^. Using the same PRISM score cutoff of 0.2, PRISM had an overall sensitivity of 0.92 and specificity of 0.95 for these species (**Fig. S2g**), and the truly-present species had significantly higher PRISM scores than when they were detected in the negative control datasets (Wilcoxon p<2e-16, **Fig. 2p**), a trend that persisted across each sequencing type individually (**Fig. S2h**). Across all the validated positive control datasets (29 taxa, 323 samples), PRISM had an overall sensitivity of 0.93 (**Fig. 2q**), and across the negative control datasets it had a specificity of 0.97 (**Fig. 2o**), demonstrating the excellent performance of PRISM using validated data.

Fifth, we tested PRISM in the scenario in which a truly-present organism was not included in its reference database. We used PRISM to analyze WGS data for 48 species (including 28 that were not in our reference database) from the CDC-HAI project^63^ in which each sample should contain only a single cultured and sequenced species. PRISM detected only one species in 29/30 samples in which the truly-present species was in the reference database (**Fig. S2i**) and predicted all detected species to be truly-present (PRISM score >0.2, **Fig. 2r**). When the truly-present species was not in the database, PRISM reported 1-5 species/sample (Wilcoxon p=1e-9, **Fig. S2i**), all of which belonged to the same genus as the truly-present taxon. Although reads assigned to these mismapped taxa continued to correspond to their mapped reference sequences (median=94% base match), they had significantly lower rates of identical base matches (Wilcoxon p<2e-16, **Fig. S2j**) and PRISM scores (median=0.03, Wilcoxon p=2.2e-14, **Fig. 2r**) compared to the truly-present taxa that were included in the reference database. PRISM classified only 4 **(4.4%)** of the 91 mismapped taxa as truly-present, demonstrating the robust performance of PRISM in this scenario. Collectively, these validation experiments demonstrate PRISM’s excellent diagnostic performance across a range of settings.

### RNA-seq based detection of microbes in CPTAC and TCGA

We next used PRISM to analyze RNA-seq data related to 8 cancer types (brain, breast, colon, head and neck, kidney, lung, ovary, pancreas) from the CPTAC (n=2075 sequencing runs) and TCGA (n=3058 runs) studies; sample details are provided in **Table S5**. PRISM initially detected between 10^0^-10^7^ microbial reads/sample and 0-200 taxa/sample across all cancer types in both CPTAC and TCGA. We used the same PRISM score of >0.2 as discussed above to select for truly-present taxa; this score was determined by examining the distribution of scores in each dataset and selecting a threshold using an elbow plot (**Fig. 3a, S3a**), where 0.2 roughly marked an inflection point in the curve. Furthermore, from the PRISM model cross-validations (**Fig. 2f**), the 0.2 threshold allowed for a model F1 score comparable to higher cutoffs while maintaining higher sensitivity (with RNA-seq data), but at the cost of lower positive predictive value. We explore the impact of choosing a more stringent PRISM score later in this report, and all data are provided in **Table S5**.

**Figure 3.**
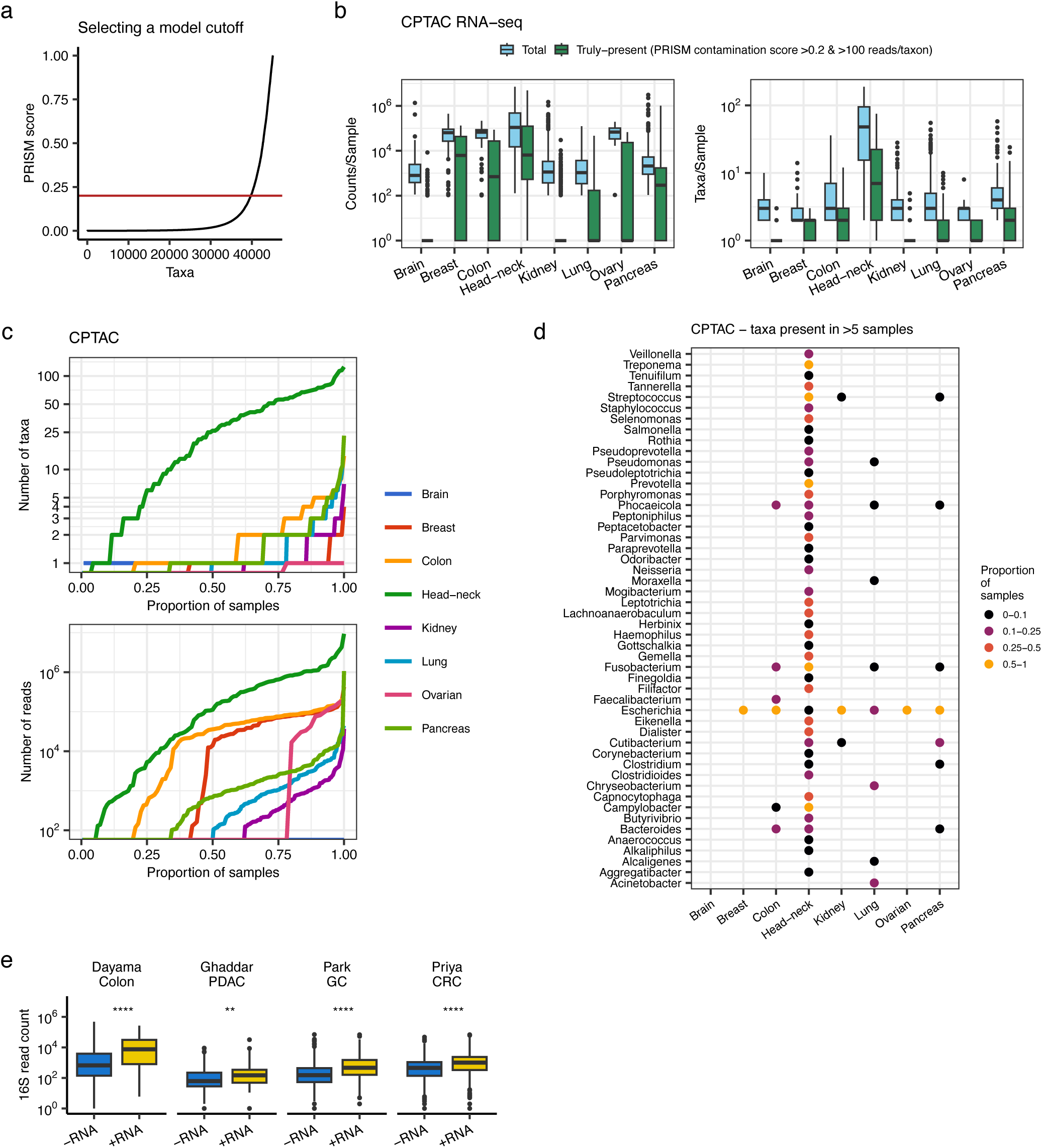
RNA-seq based detection of microbes in CPTAC. **(a)** Scatter plot of PRISM scores (0=contaminant, 1=truly-present) for all taxa identified in CPTAC. Each point is a taxon, which are ordered by their PRISM score. The red line is the cutoff used to discard likely contaminants. **(b)** Boxplots of total detected vs. truly-present microbial counts/sample (left) and taxa/sample in 8 cancer types in CPTAC. Boxplots show median (line), 25^th^ and 75^th^ percentiles (box) and 1.5xIQR (whiskers). Points represent outliers. **(c)** CPTAC samples ordered by number of microbial taxa (top) and number of microbial (reads) detected. The x-axis is scaled to reflect the proportion of samples in each cancer type. Sample curves are colored by cancer type. **(d)** Dot-plot of major genera detected in each cancer type. Dots are colored by the proportion of samples in which the genus was detected. **(e)** Boxplots of read counts in 16S metagenome sequencing of species also detected (+RNA) or not detected (-RNA) in RNA-seq performed on the same samples. Boxplots are grouped by study of origin. Boxplots are as in (b). Wilcoxon testing; **, p<0.01; ****, p<0.0001. **(f)** Bar plots indicating the number of truly-present taxa identified only by 16S metagenome sequencing (16S), only by RNA-seq (RNA), or by both in the four profiled studies. PDAC, pancreatic ductal adenocarcinoma; GC, gastric cancer; CRC, colorectal cancer.

After filtering with PRISM score of >0.2 and retaining taxa with >100 reads (the PRISM score was less reliable at <100 reads, **Fig. 2i-j**), most detected microbial content was discarded as likely contaminants or artifacts (**Fig. 3b, Fig. S3b**). Examining the distribution of taxa and reads across samples and cancer types reveals large differences in microbial load detected from these datasets. In CPTAC, only head and neck cancers had consistently widespread detectable microbes, where >75% of samples had >5 taxa and >10^3^ microbial reads detected (**Fig. 3c**). Colon and pancreatic cancer, followed by lung cancer, were next in terms of proportion of samples with substantial numbers of microbial reads and taxa detected; however, in these 3 cancer-types 75% of samples had 2 or fewer taxa detected. The other cancer types had extremely sparse microbial detection in CPTAC, i.e. >75% of samples had no microbes detected, and the remaining 25% only had 1-5 taxa detected (**Fig. 3c**).

The specific taxa detected in the CPTAC tumor samples were all ecologically relevant bacteria (**Fig. 3d, Table S5**). In head and neck tumors, PRISM identified 45 genera that were present in >5 samples, with *Fusobacterium*, *Prevotella*, *Treponema*, *Streptococcus,* and *Campylobacter* most common (present in >50% of samples); all of these taxa have previously been associated with oropharyngeal colonization and pathology^64^. In colon cancer, *Fusobacterium nucleatum*, *Bacteroides fragilis*, and *Escherichia coli* were most common and were detected at sample proportions similar to those in a targeted PCR analysis of these species in 1679 colorectal tumors^65^. Colon tumors had the highest ratio of truly-present to contaminant taxa (51%; **Table S5**). In pancreatic cancer, PRISM chiefly detected bacteria of the gastrointestinal (GI) tract. Aerobic bacteria predominated in a subset of lung tumors, although 114-12,360 reads assigned to *Phocaeicola vulgatus* were detected in 13 samples; it is uncertain whether these were true findings, or substantial contamination that passed through the PRISM algorithm*. E. coli* was detected in all cancer types except brain cancer, and it was the only bacterial species detected in breast and ovarian cancer. Although this may represent a true finding; *E. coli* was a common contaminant in the negative control datasets and one of most frequent false positives detected by PRISM (**Table S3 and Table S4**); thus, the interpretation of this finding is uncertain. Nevertheless, overall, these observations indicate that in RNA sequencing in CPTAC, head and neck tumors have abundantly detectable microbes, a subset of colon, pancreatic, and lung tumors have detectable microbes, and in the other cancer types there are sparse or no detectable microbes.

The specific taxa observations were largely not replicated in the TCGA, where >75% of all samples had 3 or fewer reported taxa (except in breast cancer) and most (∼80%) samples of the same cancer type had the same order of magnitude of reads detected (**Fig. S3c**). Such results may be interpreted as a constant level of contamination that passed through PRISM’s algorithms. Although in some cases TCGA results matched the CTPAC, such as finding *B. fragilis, F. nucleatum, and E. coli* in colon cancer and *Treponema*, *Prevotella*, *Leptotrichia*, *Fusobacterium*, and *Campylobacter* in head and neck cancer (**Fig. S3d**), many of the other taxa (e.g. *Cutibacterium, Pseudomonas, Saccharomyces, Staphylococcus*) that passed PRISM’s filters in multiple cancer types, including in breast cancer, were common contaminants in the negative control datasets (**Table S3 and Table S4**) and would be ecologically surprising to be found in these cancers. In addition, although overall there were more taxa detected per sample in TCGA (**Fig. S3e**), the taxa in TCGA (both predicted contaminants and truly-present) were detected in significantly fewer samples compared to CPTAC (**Fig. S3f**). This overall sparsity of microbial signal in the RNA-seq samples from TCGA, as well as the high prevalence of likely contaminants, limits the utility of these samples for drawing broad conclusions on the cancer microbiome. Secondly, although our experiments demonstrate that PRISM is highly effective in removing contaminants (**Fig. 2**), these TCGA results indicate that false positives may still be reported in certain datasets and thus results must be interpreted cautiously and in context of the overall findings.

### Sequencing parameters substantially impact microbial detection

The difference in RNA-seq based detection of microbes in CPTAC vs. TCGA was stark, and even in CPTAC we noted the infrequency of microbes detected in some cancer types. For example, in colon cancer we detected fewer species than in other cancer types, and in pancreatic cancer, only 10% of samples in CPTAC had >5 bacteria detected (**Fig. 3d**), whereas one recent study found bacterial content in 32% of pancreatic tumors by 16S metagenome sequencing^66^, another study found that intra-tumoral diversity could be used to stratify short and long-term survivors in two independent cohorts^21^, and a third found minimal microbial reads in pancreatic tumors^22^. We hypothesized that some of these differences between CPTAC, TCGA, and the other studies were in part due to differences in sequencing approach, the impact of which would be exacerbated in low biomass settings.

To address this hypothesis, first we examined the impact of the RNA-seq selection strategy on microbial detection, as CPTAC3 used ribosome-depletion (RD) and CPTAC2 (breast, colon) and TCGA used poly-A selection (PAS), which biases against microbial reads since microbes lack poly-A modifications. We had already observed in CLID that the PRISM score was less sensitive in PAS vs. RD data (**Fig. 2h**). However, irrespective of the PRISM score (as cutoffs can be subjective), we asked whether RD captured more total microbial content. To address this point, we examined the CLID contaminants, on the presumption that those samples had similar contamination levels since they were from controlled cell-line experiments intended to study host responses to one or a few introduced pathogens. We found that (1) the RD strategy captured a significantly greater number of taxa per sample than PAS (Wilcoxon, p=4.1e-7, **Fig. S3g**), (2) taxa in the RD data were detected in a significantly higher proportion of samples (Wilcoxon, p<2e-16; **Fig. S3h**) and (3) at nearly twice the relative abundance compared to PAS samples (Wilcoxon p<2e-16; **Fig. S3i**), and (4) the RD samples had a significantly higher taxa read counts Shannon diversity (Wilcoxon p<2e-16; **Fig. S3j**). These observations indicate that for RNA-seq, RD is overall more reliable than PAS at capturing microbial reads. These findings suggest one mechanism leading to more consistent microbial detection across samples in CTPAC3 compared to TCGA and as well as the unexpected lack of abundant microbes in CPTAC2 samples (breast and colon tumors).

Second, we examined the extent to which batch effects influenced the CPTAC and TCGA findings. For CPTAC, sequencing was done per project, however the RNA selection varied between CPTAC2 (PAS; breast, colon cancer) and CPTAC3 (RD; all others). Similar to the finding above, the samples with PAS had fewer taxa detected (**Fig. 3d**). In TCGA, analyses were done at multiple sequencing centers. For colon and head and neck cancers in TCGA where multiple taxa were detected in concordance with CPTAC, we visualized the samples on a principal component plot computed from taxon relative abundances. Samples of the same cancer type were more similar to each other, and samples did not cluster by sequencing center, indicating that PRISM was identifying signal independent of this batch variable (**Fig. S3k**). However, it is possible for batch effects to bias both truly-present and contaminant signals. PRISM does not inherently overcome such effects, but identifies the most robustly supported taxa within the data. Considering the possibility of batch effects is necessary.

Lastly, we suspected that other tumors in CTPAC and TCGA may have had microbes that bulk RNA-seq was not able to capture. To investigate the sensitivity of RNA-seq of human samples to capture low biomass microbiomes, we analyzed data by PRISM from 4 studies (pancreatic cancer^32^, colorectal cancer^67^, gastric cancer^68^, and colonic tissue without detectable malignancy^69^) that performed both RNA-seq and 16S metagenome sequencing on the same tissues. Our hypothesis was that species detected by RNA-seq (irrespective of their PRISM score) were at higher abundance than species not detected and thus would have higher read counts in the 16S data. In 159 samples from the 4 studies, using PRISM we confirmed that RNA-seq was less sensitive to capturing microbial sequences compared to metagenome sequencing. Species detected in RNA-seq had significantly higher 16S read counts than those not detected in all 4 studies (Wilcoxon all p<0.003; **Fig. 3e, Table S6**). To confirm that the species that were not detected in RNA-seq were not captured (as opposed to missed by PRISM), we checked for their read counts in the RNA-seq samples using the highly sensitive Kraken2 method and found a median of zero reads for those species. These data indicate that lack of microbial detection in human RNA-seq cannot always be presumed to show absence.

### Microbial associations with pancreatic cancer in CPTAC

Despite the above limitations in sensitivity of RNA-sequencing of human tissues to detect microbes, PRISM detected microbes in many CPTAC samples. Therefore, the presence of associated data in CPTAC provides an opportunity to investigate microbial associations with molecular and clinical attributes of these tumors. We focused on pancreatic cancer, where there has been recent debate about microbial influence^21,22,32^, first examining the species identified in two or more tumors with >100 reads. We identified *E. coli* in 66 tumors as well as 29 other species, each detected in 2-7 tumors; most species were known oral or gut bacteria (**Fig. 4a**). Most detected microbial products were related to ribosomal RNA, accounting for 89% of the products (**Fig. 4b**); this was an expected finding, increasing confidence in our analytical approach.

**Figure 4.**
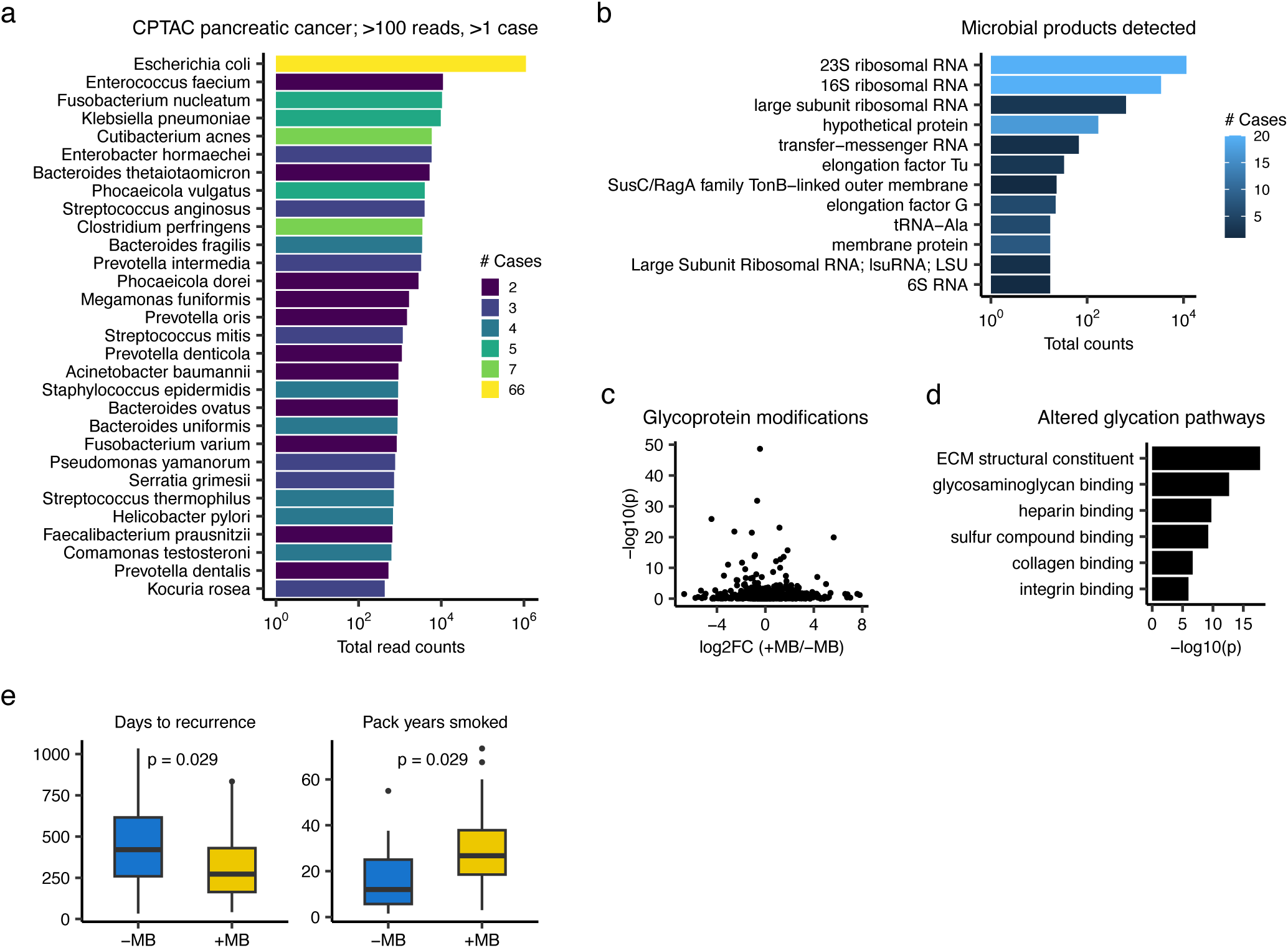
Molecular and clinical associations with the pancreatic cancer microbiome. **(a)** Bar plot indicating the total read counts of species identified in 159 cases of pancreatic cancer in CPTAC. Bars are colored by the number of cases in which the species was detected. **(b)** Bar plot of the most common microbial products detected in CPTAC pancreatic cancer. **(c)** Volcano plot of genes with glycoprotein modification. The x-axis is the log2 fold change of average gene modification level in tumors with detectable microbiome (+MB; n=91) vs. those without (-MB; n=68). Y-axis indicates the Wilcoxon p-value. **(d)** Bar plot indicating gene ontology pathways of the genes with significantly different glycoprotein modifications (p<1e-5) in tumors with vs. without detectable microbiome. (e) Boxplots of the days to tumor recurrence (left) and pack years smoked (right) of pancreatic cancer patients whose tumors have (+MB) vs. do not have (-MB) detectable microbes using RNA-seq. Boxplots show median (line), 25^th^ and 75^th^ percentiles (box) and 1.5xIQR (whiskers). Points represent outliers. P-values are from Wilcoxon testing.

Then, using the species cutoffs of >100 reads, detection in two or more tumors, and PRISM score >0.2, we identified 88 tumors with detected microbes and 71 without in the CPTAC pancreatic cancer cohort. We next asked whether tumors differing in whether microbes were detected also differed in their molecular characteristics. Using Wilcoxon testing and multiple hypothesis correction, we found no significant association between microbial detection in tumors and number of gene mutations or levels of mRNA expression, protein expression, protein phosphorylation, or protein methylation. However, there was a significant difference in their glycoproteomic properties; we identified 58 genes with significant differences in mean gene modification level between tumors with microbe-detected vs. absent (Wilcoxon adjusted p-value < 0.05, **Fig. 4c**). Since most of these genes had small average log-fold differences between the groups, we next conducted gene ontology (GO) analysis of the significantly altered glycated proteins. GO analysis identified several significant pathways that were related to extracellular matrix remodeling (**Fig. 4d, Table S7**). When we repeated GO analysis with shuffling the glycated gene name labels, no significant glycoprotein differences were found between tumors with detected microbes vs. absent, indicating that chance alone did not explain our results. Lastly, we asked whether there were any clinical attributes linked to the detection of microbes (**Fig. 4e**); patients whose tumors had detectable microbes had significantly fewer days to tumor recurrence (Wilcoxon p=0.029 and significantly more substantial smoking history (pack-years smoked; Wilcoxon p=0.029). We also repeated the clinical association testing removing *E. coli*, as it could represent a biological contaminant that passed through PRISM’s filters, and the associations remained consistent (**Fig. S4a**).

Finally, to assess the robustness of these findings, we repeated these association tests for the CPTAC pancreatic cancer samples using the more stringent PRISM score cutoff of >0.4. This identified 65 tumors with detected microbes and 94 tumors without. We identified the bacterial species (**Fig. S4b**) and products (**Fig. S4c**) and tested for tumor microbial association with glycoprotein modifications (**Fig. 4d-e**) and clinical attributes (**Fig. S4f**). All of the results remained consistent, with the exception of partial differences in the glycoprotein pathways identified. Taken together, these results indicate that a detectable difference in microbial content in pancreatic tumors is associated with specific cellular activities and clinical parameters, despite the limitations of RNA-seq to capture low-biomass microbiome.

## Discussion

In developing PRISM, we documented four major sources of false-positive detection of microbes in specimens with low microbial biomass and we designed approaches to mitigate them. The first source of error is human reads incorrectly classified as microbial, many that have been missed by multiple algorithms. Thorough human read removal is crucial, including checking for alignment to sequences that are not part of a standard human reference genome including alternative haplotypes, repetitive mitochondrial and ribosomal sequences, and unplaced or unlocalized scaffolds^70^. PRISM does this efficiently using Kraken, STAR, and BLAST on a taxa-representative subsample of data. The second source is incorrect taxon assignment of k-mer-based mapping, where assigned reads have no or poor full-length read mapping, which PRISM identifies in its BLAST analysis. The third source of error is multi-mappable microbial reads. We found that many of the initially reported taxa by Kraken2 had no reads that uniquely mapped to them, i.e. they were attributable only to reads that could also be mapped to other taxa. Focusing only on taxa with uniquely mapping reads always identified the known truly-present taxa in our datasets and reduced the burden of false-positive taxa. The fourth source is contamination. Using our large training datasets, we identified patterns of microbial sequence capture and implemented a machine learning model to compute a PRISM score (i.e. the probability of true presence and not a contaminant) for each taxon in a sample. This score adds a measurable parameter to prioritize taxa that are highly likely to have originated from the source tissue. However, in practice, score cutoffs can be selected in congruence with taxon read counts and PRISM score distributions in a sample. While not error-free, at the least, PRISM substantially reduces technical artifacts that are exacerbated in the low-biomass setting and reports the most uniquely identifiable taxa. The second layer provided by the PRISM score further prioritizes taxa and identifies whether a signal in a dataset should be pursued.

While PRISM substantially reduces false positives and identifies high-confidence microbes, our experiments showed that false negatives also represent a major source of error. Our analysis of 16S metagenomic sequencing vs. human bulk RNA-seq based on common samples showed that in low biomass settings, metagenomics is more sensitive, and that microbes of lower abundance in the 16S data were less likely to be captured by bulk RNA-seq. Interpreting the absence of microbial data in non-metagenomic (e.g. mammalian) sequencing thus also requires caution; targeted and high depth sequencing coupled with orthogonal validations (e.g. quantitative PCR, fluorescence in situ hybridization, immunohistochemistry) and biological plausibility are necessary for definitive studies.

After validating PRISM’s performance, we used it to detect microbes in RNA-seq data of 8 cancer types in the CPTAC and TCGA datasets. In CPTAC, head & neck and gastrointestinal tract tumors harbored rich microbiomes, a subset of pancreas and lung tumors had detectable microbes, and there was little evidence for widespread tumor microbiome in the other cancer types. Microbiome data in TCGA was more sparse which limited its analysis and the ability to draw conclusions, possibly due to insufficient microbial capture in part due to poly-A selection. In CPTAC, for pancreatic cancer, we found that the presence of detectable microbes was associated with altered protein glycosylation, consistent with evidence linking glycosylation changes to altered host-microbiome interactions and immune dysregulation in inflammatory bowel disease and some cancers^71,72^. Whether these microbes alter the tumor glycome or an altered glycome attracts microbes is an important open question. We also found that microbial presence was associated with a more substantial smoking history and increased risk of tumor recurrence in pancreatic cancer. Whether microbes are an active facilitator of tumor recurrence or a consequence of it is another important area for future investigation. Overall, our findings indicate that microbes exist in at least a subset of tumors and cancer types and their presence may be associated with altered molecular and clinical attributes. However, since their abundances are generally low, they are best assessed through targeted metagenomic approaches. Human sequencing datasets can be useful for hypothesis generation, but cross-study comparisons may be limited, and negative data should be interpreted with caution as there is an unknown lower threshold of detection. As such, in this manuscript, we report microbes detectable with high-confidence in these datasets and we do not attempt to define definitive cancer microbiomes.

Our study has several strengths and limitations. The first strength is the uniquely large datasets assembled for the development of PRISM and its realistic and challenging benchmarking experiments; these provide confidence in our methods. Second, PRISM directly interrogates reads via both fast k-mer profiling and full-length mapping, addressing root issues of mapping uncertainties. Third, PRISM can be used for any sequencing approach, including single-cell, bulk, metagenomic, and RNA or DNA sequencing and is highly computationally efficient. We also identified seven possible limitations. First, there is no objective method to select a PRISM score cutoff, and the use of differing scores can alter interpretation of results. We advise selecting PRISM scores in the context of other parameters, such as taxa counts, distribution of PRISM scores in a sample or study, and other factors such as the number of samples in which a taxon was detected. Second, PRISM conservatively identifies uniquely identifiable taxa, but could fail to identify highly similar strains, although we did not observe this in our testing experiments. Third, PRISM’s BLAST steps perform competitive mapping using pre-identified taxa from Kraken2. The final output is highly dependent on these taxa; if a taxon is missed by Kraken2, or if it reports a large number of highly similar taxa, the truly-present taxon will be harder to identify. Fourth, PRISM’s results will be limited by errors in reference genomes used and when known model organism genomes are not included. Fifth, PRISM relies on patterns of microbial sequence capture observed in our training and testing datasets, many of which were cell line experiments or sequencing of microbial isolates; these patterns may differ from those of complex microbial communities in human tissues. Sixth, PRISM removes reads that map to human sequences using BLAST, some of which are labeled as “chimeric clones”; this conservative approach risks removing reads of true microbial origin. Seventh, the PRISM score favors a higher diversity of reads; it may be artificially low for species that are dormant, more transcriptionally limited, have lower sequence variation, or are less characterized in reference databases.

In conclusion, PRISM facilitates the correct ascertainment of microbes, especially in low-biomass settings. Despite limitations that are intrinsic to detecting rare phenomena, PRISM provides new information identifying microbial presence in certain human malignancies and their absence in others.

## Methods

### PRISM algorithm

PRISM identifies truly-present microbial species from genomic sequencing data and substantially reduces the burden of false-positive taxa due to missed human reads, errors of k-mer based taxonomic assignment, multimapping microbial reads, and contamination. To do this, it has six key parts: (1) surveilling genomic data with Kraken2, (2) creating a microbial taxa-representative subsample of data, (3) identifying and removing missed human reads in the subsampled data with STAR, (4) full-length alignment of the subsampled data with BLAST and determining the uniquely identifiable taxa, (5) machine learning-based PRISM score prediction, and (6) a final BLAST of all data against the identified taxa to report final taxa and gene counts. The ordering of these steps was designed for computational efficiency – Kraken rapidly removes most human reads, STAR is faster than BLAST and catches many missed reads, and BLAST finds the remaining reads. This is faster than running a human alignment step first on the full file; however, in general, mapping first to the human genome and removing those reads will increase PRISM’s speed and will reduce the number of false-positive microbes initially reported by Kraken.

Importantly, barcode and index sequences should be removed prior to analysis with PRISM as this can affect mapping statistics. All algorithmic steps are described in further detail below.

*(1). Initial surveillance with Kraken2*. This first step processes sequencing data using Kraken2^33^, a sensitive k-mer based taxonomic classifier, with a reference database of complete microbial (bacterial, fungal, viral) genomes, the human genome, and known vector sequences recorded in RefSeq^34^ as of August 2024 using default Kraken2 parameters. This allows fast removal of the majority of human reads and vector sequences and identifies the possible microbial taxa present in the sample.

*(2). Creating a taxa-representative subsample of the data*. After extracting the Kraken2-identified putative microbial reads, PRISM randomly samples up to *n* (default=1000) reads from each taxon and creates a subsampled dataset for further interrogation. Taxa with >*m* reads (default 10 reads) are subsampled. If a taxon has <*m* reads directly assigned to it by Kraken2, reads assigned to its clade (i.e. subspecies or strains) are sampled, or if none exist, reads assigned to higher levels of the taxon’s phylogeny are subsampled. The result is a small, representative dataset that can be efficiently interrogated in downstream steps.

*(3). Removing human reads with STAR*. Next, PRISM aligns the subsampled data to the human genome using STAR^35^ with default parameters. Human mappable reads are removed from the subsampled dataset and the remaining microbial taxa are identified. In this manuscript, we used human genome version hg38. This step is only applicable for RNA-seq datasets; DNA-based datasets still benefit for a second line of human-read removal in the next BLAST steps.

*(4). Aligning the subsampled data with BLAST.* Next, PRISM aligns the subsampled data to human genomic sequences (NCBI taxid:9606), genomes of common model organisms, and the remaining microbial genomes using default BLAST parameters. This step performs competitive mapping and allows PRISM to identify missed human reads that are not part of a standard genome draft, such as repetitive ribosomal sequences. It also identifies not uncommon genomic material of common model organisms. In this manuscript, we included all taxonomy IDs of all species of the *Rattus and Mus* genera, although users can specify other species. PRISM retains only those reads that mapped to microbial species and determines which taxa are uniquely identifiable by identifying the taxa of reads that mapped to only one taxon. PRISM then prioritizes the BLAST results to assign reads to these uniquely identifiable taxa.

*(5). PRISM score prediction.* For each remaining taxon in the subsampled data, PRISM, next computes a set of features that are used by PRISM’s pretrained xgboost^73^ model to predict a PRISM score for that taxon. The score represents the probability that a taxon is a contaminant (0) or is truly-present (1). The PRISM model uses 6 classes of features for a total of 55 variables, as next described.

The first class of features quantifies taxon gene-product diversity using the BLAST results and is based on the observation that truly-present species express a higher diversity of genes compared to contaminants. These include “fprod”, the proportion of BLAST products in the sample assigned to the taxon; “fuprod”, the proportion of unique products in the sample assigned to the taxon; “fugene”, the taxon’s proportion of unique genes in the sample; “prod_div”, the Shannon diversity of the taxon’s product counts; “gene_div”, the Shannon diversity of the taxon’s gene counts. Shannon diversity is calculated as SD = -∑*p_i_*ln(*p_i_*), where *p_i_* is the proportion of genes or products attributed to gene or product *i.* Because the Shannon diversity correlates with the total number of observations, PRISM does 10 iterations of randomly sampling 100 genes or products and calculating the Shannon diversity, then averages the 10 diversity measures.

The second class of features relates to read multi-mappability and is based on the observation that truly-present species have more uniquely mapping sequences than contaminants. These features are: “ratio”, the proportion of taxon’s reads that are multimappable; “sacc_ratio4”, the ratio of the number of accessions only mapped by uniquely mapping reads to the number of accessions mapped by both uniquely and multi-mapping reads; “sacc_ratio5”, the ratio of the number of accessions only mapped by multimapping mapping reads to the number of accession mapped by both uniquely and multi-mapping reads.

The third class of features is based on Kraken2’s read count estimates and is based on the observation that truly-present species have more total and unique reads than contaminants, as estimated by Kraken2. The features are: “n2”, the total reads assigned to the clade rooted at the taxon; “n3”, number of reads specifically assigned to the taxon; “uniq”, estimated number of unique k-mers assigned to the taxon; “fmicro”, taxon read counts relative to the total reads counts assigned by Kraken to bacteria, fungi, and viruses.

The fourth class of features is “k-mer phylogeny”, or the proportion of k-mers per taxon assigned to each taxonomic level by Kraken2. These features are based on the observation that truly-present species have more reads assigned to the species level than contaminants. Levels for which k-mer phylogeny proportions are computed include kingdom, phylum, class, order, family, genus, species, unclassified, and host (human).

The fifth class of features relates to misclassifications by Kraken2 and is based on the observation that Kraken2 assigns more reads to taxa related to truly-present species than contaminants. For the kingdom (k_), phylum (p_), class (c_), order (o_), family (f_), genus (g_), and species (s_) levels, PRISM determines the proportion of reads (_r), unique k-mers (_u), and species (_n) in the sample that are assigned by Kraken2 to the phylogenetic lineage of each taxon.

*(6). Final BLAST*. After determining the uniquely identifiable taxa and computing a PRISM score for each, PRISM identifies reads that could be attributable to these taxa by examining the previous Kraken2, STAR, and BLAST results. It extracts these reads and BLASTs them to the uniquely identifiable taxa list. PRISM then merges these BLAST results with GenBank^74^ gene and product data and reports final taxa counts, PRISM scores, and gene and product data, summarized at each taxonomic level.

### Assembling the PRISM datasets, training the contamination model, and benchmarking

The cell line infection dataset (CLID) was used for the initial development of PRISM. These were RNA-seq experiments of human cell-lines that had been purposefully infected with a known pathogen. These datasets were identified by querying the Sequence Read Archive for “((((“public”[Access]) AND “rna seq”[Strategy]) AND “transcriptomic”[Source]) AND cell line) AND “Homo sapiens”[orgn: txid9606]” and searching for specific microbial species names. All samples were manually verified that they met our desired criteria. In each CLID sample, the known pathogen(s) were truly-present (each sample had 1-4 known pathogens), and all other taxa detected were contaminants. These datasets are listed in **Tables S1** and **S3**.

To train PRISM’s contamination model, we combined CLID with 4 other datasets that we generated which had a larger number of known truly-present species. The first dataset consisted of synthetic combinations of CLID which we called CLID-C. For each CLID-C sample, between 100-1000 known pathogen reads from different CLID samples were randomly sampled and combined with reads from known CLID contaminants. In total, we created 153 new CLID-C samples; each sample had between 4-45 truly-present species and between 1-201 contaminants, all from RNA-seq data. To prevent data leak during model cross-validation, we created CLID-C samples in 5 folds, such that there was no crossover of data from SRA projects amongst folds, i.e. all data from the same SRA project were in the same fold. These samples are denoted by projects m1-m5 in **Table S3**. Details of their sample origins and taxa are in **Table S8**.

We also created a dataset called WGS which contained *in silico* combinations of cultured and whole genome sequenced bacterial isolates from human gut^36^, bladder^37^, and the FDA ARGOS database^38^. For these data, from each original sample we randomly sampled reads assigned to the cultured species by Kraken2. We then randomly combined these into 236 new WGS-C samples, each with 1-104 truly-present species. Samples were created in 5-folds, with 2-folds each for the gut and bladder data and one-fold for the FDA ARGOS data, with no crossover between samples across the folds. These samples are denoted by projects f1-f5 in **Table S3**. Details of their sample origins and taxa are in **Table S8**. The WGS dataset also included negative control WGS data for n=138 samples; these were reagent or blank controls from whole metagenome sequencing studies^39–44^.

Lastly, we added data from metatranscriptomic studies from human gut^45^, mouth^46^, skin^55^, and vagina^47^, which we called META. To cautiously avoid including contaminants from these datasets, we used Kraken2 to identify and extract reads from taxa with reads greater than the median reads/taxa in its sample. These samples were grouped into 4-folds, one for each study. The combined dataset of CLID, CLID-C, WGS-C, and META contained 34,464 total observed taxa, 630 unique true-positive taxa, 2,172 unique contaminants, 19,420 total contaminant taxa, and 15,044 total true-positive taxa. Sample details are in **Table S3**.

We explored these data extensively and engineered the features discussed in the PRISM algorithm section. These were created heuristically as we searched for metrics that distinguished truly-present and contaminant taxa. We computed those features for our dataset and then used xgboost^73^ to train a gradient-boosted decision tree model to classify truly-present vs. contaminant taxa. The model was evaluated using 5-fold cross-validation and the predesigned folds discussed above. The model used the following parameters: nrounds = 100, reg_lambda = 10, colsample_bynode=0.25, subsample=0.5, eval_metric = ’auc’, objective = ’binary:logistic’. Although these parameters were chosen, they did not have a substantial impact on the final results over other parameter choices (data not shown).

We benchmarked PRISM’s cross-validation performance on the CLID dataset against three other high sensitivity or specificity methods for taxonomic classification (Kraken2^33^, MetaPhlAn^48^, Metabuli^49^) and one method for computational decontamination (SAHMI^31^). All methods were run with default parameters (for SAHMI, we used a negative control percentile cutoff of 0.95). We included an additional step of filtering a “blacklist” of 497 common contaminating genera^17^ from the output of these methods (but not from PRISM). Doing this did not reduce sensitivity of any method but slightly increased their specificity.

### CPTAC and TCGA analyses

All fastq files and clinical metadata were obtained directly from the Genomic Data Commons^75^, and sequencing reads unmapped to the human reference were extracted from the BAM files for analysis with PRISM. From CPTAC, we downloaded RNA-seq data of tumor tissues from brain, breast, colon, head & neck, lung, ovary, and pancreas cancers (CPTAC3 and CPTAC2 datasets; endometrial cancer was not analyzed). From TCGA, we downloaded RNA-seq data of tumor tissues from glioblastoma (GBM), breast cancer (BRCA), colon adenocarcinoma (COAD), head and neck squamous cell carcinoma (HNSCC), lung adenocarcinoma (LUAD), ovarian cancer (OV), and pancreatic ductal adenocarcinoma (PAAD). Molecular (RNA-seq, mutation, proteomic, phosphoproteomic, glycoproteomic) data from CPTAC pancreatic cancer were obtained from the relevant publication^76^. All data were processed using the standard PRISM pipeline. In this manuscript, we used a PRISM score cutoff of >0.2, we filtered for microbial reads with >80% sequencing matching (pident) in BLAST, and we retained taxa with >100 reads. We considered several variants to these parameters, and all raw taxa count data are reported in **Table S5**. For the pancreatic cancer microbiome association testing with molecular data, we did Wilcoxon testing with individual gene mRNA, protein, phosphoprotein, and glycoprotein levels and with total number of tumor mutations. The gene ontology analysis for the significantly altered glycoproteins was done using the clusterProfiler^77^ R package with default parameters; the full human proteome was used as the gene background for this analysis. When evaluating batch effects in TCGA, we constructed principal component plots for colon and head and neck cancer using taxa relative abundances in samples. We included data for all taxa detected in these samples and normalized data per sample by log_10_(TP10k+1).

### Statistical analyses

All statistical analyses were performed using R version 4.3.1. All p-values were corrected for false-discovery rate (fdr) for multiple hypotheses using the p.adjust function with method= “fdr”, unless otherwise stated. The tidyverse package was substantially used for data processing (https://www.tidyverse.org/). The ggpubr package (https://github.com/kassambara/ggpubr) was used to compare group means with nonparametric tests and to perform multiple hypothesis correction for statistics that are noted in the figures. P-values reported as <2×10^-16^ result from reaching the calculation limit for the native R statistical test functions and indicate values below this number, not a range of values.

## Data availability

All datasets analyzed in this study are listed in Tables S3-S6. All datasets generated in this study are available on the PRISM Github: https://github.com/sjdlabgroup/PRISM and from the authors upon reasonable request.

## Code Availability

The PRISM pipeline is available on our Github: https://github.com/sjdlabgroup/PRISM

## Acknowledgments

We acknowledge the Office of Advanced Research Computing (OARC) at Rutgers, The State University of New Jersey for providing access to the Amarel cluster URL: https://it.rutgers.edu/oarc. We also acknowledge grant support from National Institutes of Health grant R35GM149224 (SD), National Institutes of Health grant U01AI22285 (MJB), Sergei Zlinkoff Foundation (MJB), Canadian Institute for Advanced Research (MJB).

## Author contributions

BCG conceived the study and designed and performed all analyses. BCG, MJB, SD interpreted the data and wrote and revised the manuscript.

## Figure legends

**Figure S1.**
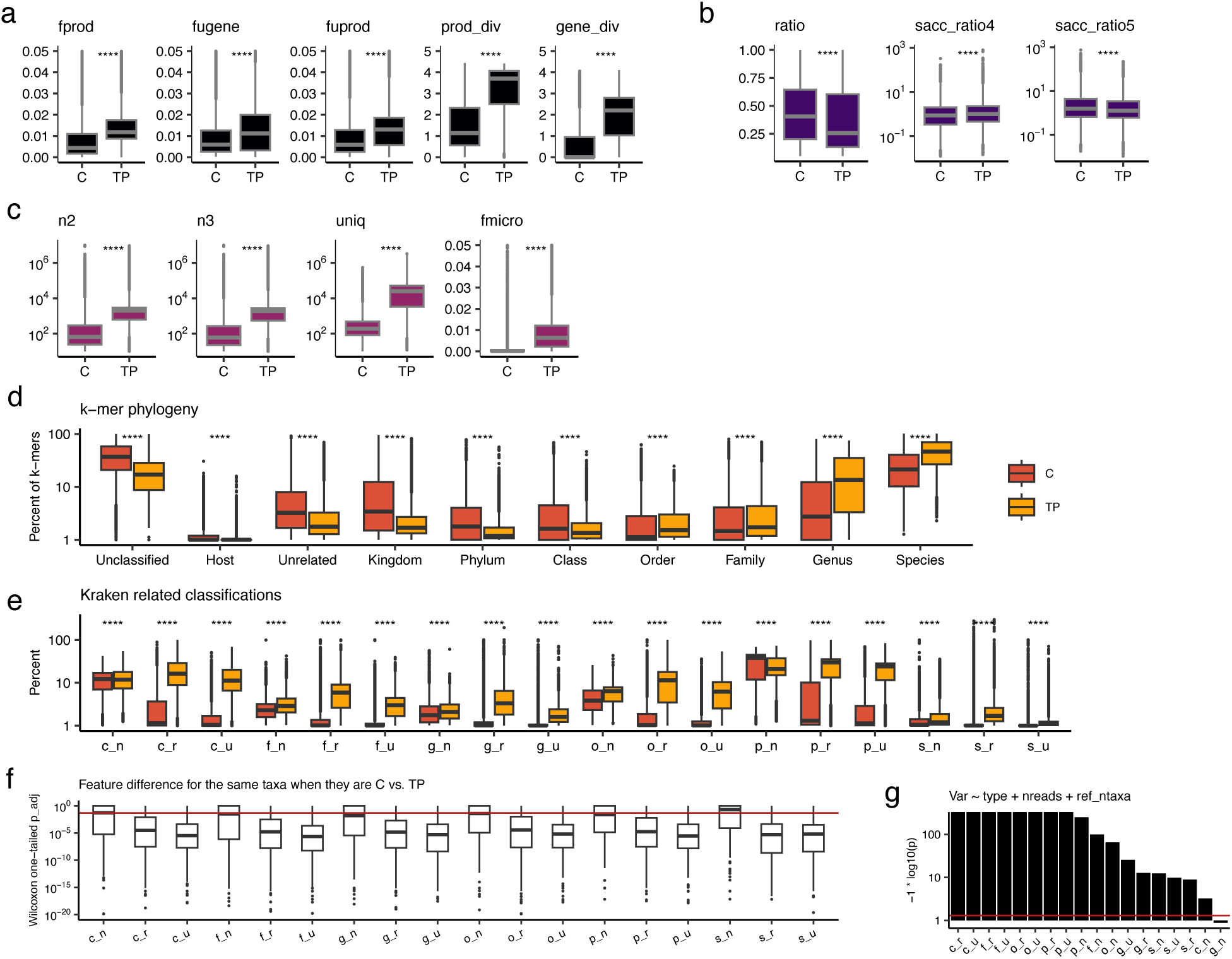
PRISM contamination model features. Boxplots comparing the values of PRISM model features between known contaminant (C) taxa and truly-present (TP) taxa in the CLID datasets. See Methods for specific feature details and formulas. In all panels, boxplots show median (line), 25^th^ and 75^th^ percentiles (box) and 1.5xIQR (whiskers), and points represent outliers. Wilcoxon testing; ****, p<0.0001. **(a)** Features related to microbial and gene and product (as reported by BLAST) statistics calculated from BLAST output; fprod, proportion of total products; fuprod, proportion of unique products; fugene, proportion of unique genes; prod_div, products Shannon diversity; gene_div, genes Shannon diversity. **(b)** Statistics per taxon related to the proportion of reads that are uniquely vs. non-uniquely mappable. mb_strand, strandedness (right vs. left) of mapped reads; mb_uniq, number of unique BLAST accession IDs mapped. **(c)** Statistics per taxon calculated from the Kraken2 report. n2, total reads assigned to the clade rooted at the taxon; n3, number of reads specifically assigned to the taxon; uniq, estimated number of unique k-mers; fmicro, taxon read counts relative to the total reads counts assigned by Kraken to bacteria, fungi, and viruses. **(d)** Proportion of k-mers per taxon that Kraken2 assigned to each phylogenetic class. **(e)** For each taxon, the number of reads (_n), unique k-mers (_u), or species (_n) that Kraken2 assigned to the same phylogenetic lineage for each level of the phylogenetic tree. (**f**) Boxplots show one-tailed Wilcoxon p-values testing for a difference in Kraken2-related classification feature values for the same taxa when they were contaminants vs. truly-present. P-values were adjusted for false-discovery. The red line marks p=0.05. (**g**) We used multiple linear regression models to test for the significance of the Kraken-related classification features while controlling for taxon read abundance and the number of related taxa (belonging to the same class) present in the reference database. Bars indicate the coefficient p-value for the tested feature. The model tested feature ∼ taxon type (contaminant vs. truly present) + read count + number of same class taxa in the reference database. The red line indicates p=0.05.

**Figure S2.**
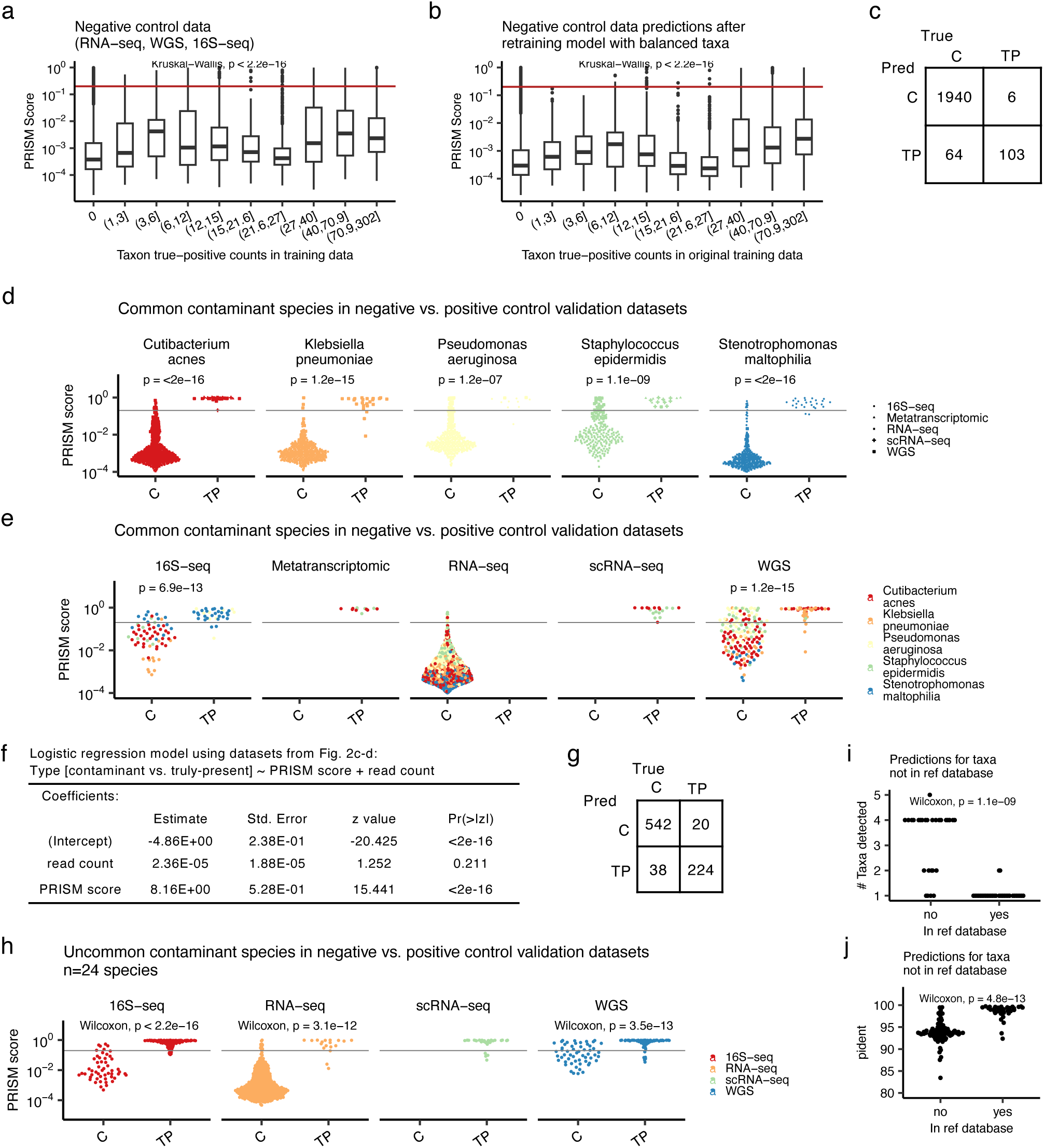
Validating the PRISM score. **(a)** Boxplots of PRISM scores for the RNA-seq data from cell lines in the absence of known infection, a negative control validation dataset to test for model prediction bias toward over-represented true-positive taxa in its training data. Data are stratified into deciles related to the number of instances in the training data for each true-positive taxon. The red line marks 0.2, the true-positive threshold used in this paper. Boxplots show median (line), 25^th^ and 75^th^ percentiles (box) and 1.5xIQR (whiskers), and points represent outliers. (**b**) Repeating the analysis from (a) but after subsampling the training data to include 15 instances of each contaminant or each truly-present taxon. The new model is then used to predict PRISM scores for the negative control dataset. Taxa are stratified into the same deciles as in (a). Boxplots are as in (a). (**c**) Confusion matrix of PRISM predictions for the data from Fig. 2p. Data are for five species that were common known contaminants (C) in the negative control datasets vs. when they were validated as truly-present (TP) in human infections. (**d**) Swarm plots of PRISM scores for the data from Fig. 2p, stratified by species; p-values indicated Wilcoxon testing. (**e**) Swarm plots of PRISM scores for the data from Fig. 2p, stratified by sequencing technology; p-values indicated Wilcoxon testing. (**f**) Logistic regression modeling of the data from Fig. 2p assessing the significance of the PRISM score vs. read count in distinguishing contaminant vs. truly-present species. (**g**) Confusion matrix of PRISM predictions for the data from Fig. 2o. Data are for 24 species that were uncommon contaminants (C) in the negative control datasets vs. when they were validated as truly-present (TP) in human infections. (**h**) Swarm plots of PRISM scores for the data from Fig. 2o, stratified by sequencing type; p-values indicated Wilcoxon testing. (**i**) Swarm plot of the number of taxa detected in CDC-HAI WGS of human pathogens for taxa that were (yes, n=20) or were not (no, n=28) in our reference database. (**j**) Swarm plots of percent identical base match (pident) from BLAST for taxa that were or were not in the reference database.

**Figure S3.**
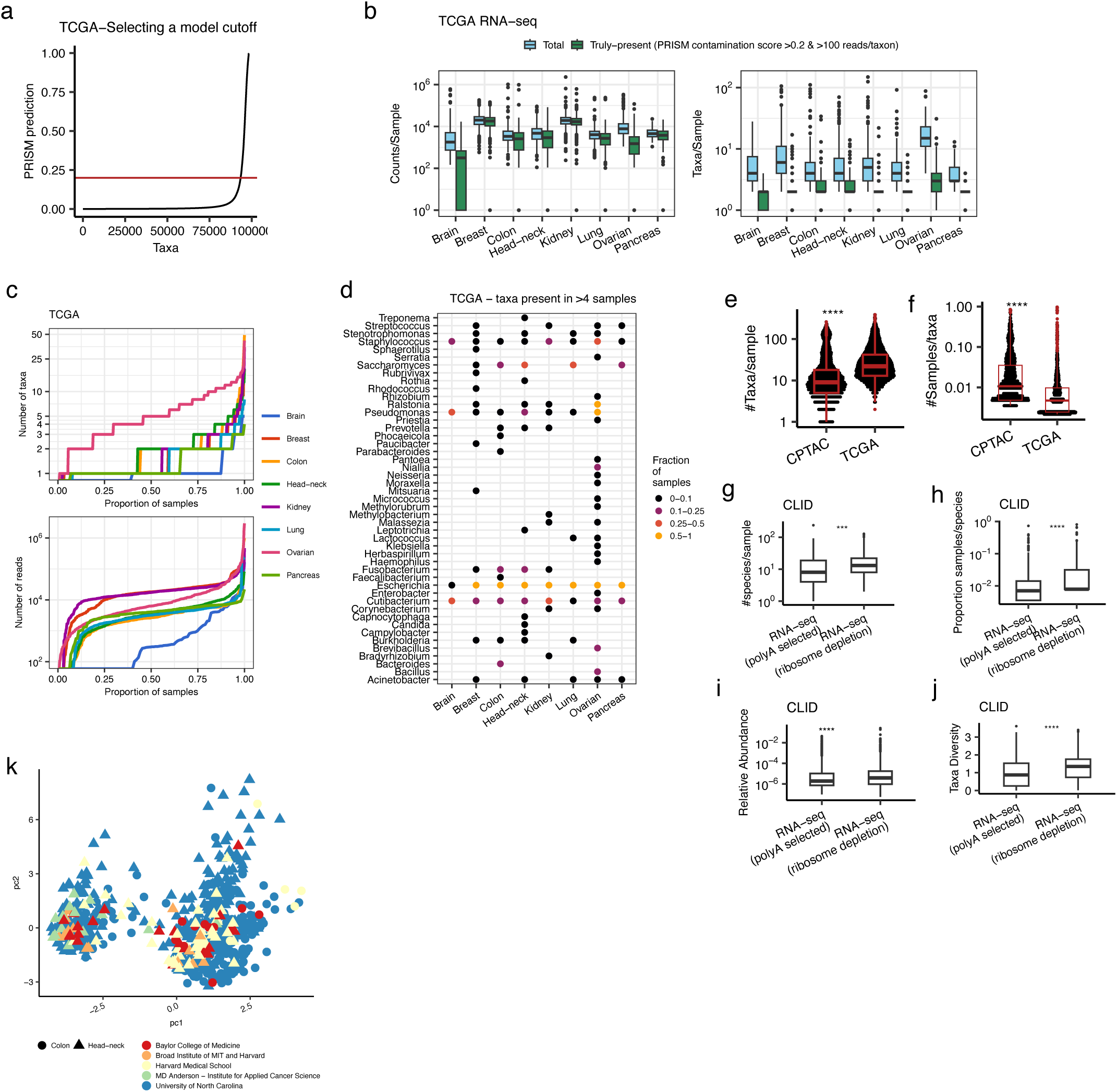
Profiling the microbiome in TCGA. **(a)** Scatter plot of PRISM scores (0=contaminant, 1=truly-present) for all taxa identified in TCGA. Each point is a taxon, which are ordered by their PRISM score. The red line is the cutoff (PRISM score > 0.2) used to discard likely contaminants. **(b)** Boxplots of total detected vs. truly-present microbial counts/sample (left) and taxa/sample in 8 cancer types in TCGA. Boxplots show median (line), 25^th^ and 75^th^ percentiles (box) and 1.5xIQR (whiskers), and points represent outliers. **(c)** CPTAC samples ordered by number of microbial taxa (top) and number of microbial (reads) detected. The X-axis is scaled to reflect proportion of samples in each cancer type. Sample curves are colored by cancer type. **(d)** Dot-plot of major genera detected in each cancer type. Dots are colored by the proportion of samples in which the genus was detected. **(e)** Boxplots and swarm plots of the number of taxa/sample detected in CPTAC and TCGA. (**f**) Boxplots and swarm plots of the proportion of samples that each taxon was detected in for taxa detected in CPTAC and TCGA. (**g**) Boxplots showing the number of species per sample detected in CLID in poly-A selected (PAS) vs. ribosomal-depletion (RD) RNA-seq. (**h**) Boxplots comparing the proportion of samples in which each taxon in CLID was detected in PAS vs. RD samples. (**i**) Boxplots comparing the relative abundance of contaminants in CLID detected in PAS vs. RD samples. Contaminants are presumed to be present at a similar level of total abundance in these samples. (**j**) Boxplots comparing the diversity of taxon read counts in CLID in PAS vs. RD samples. (**k**) Principal component plot of TCGA colon and head and neck cancers computed based on taxa relative abundances. Each point represents a sample. Points are colored by their sequencing center and shaped by cancer-type.

**Figure S4.**
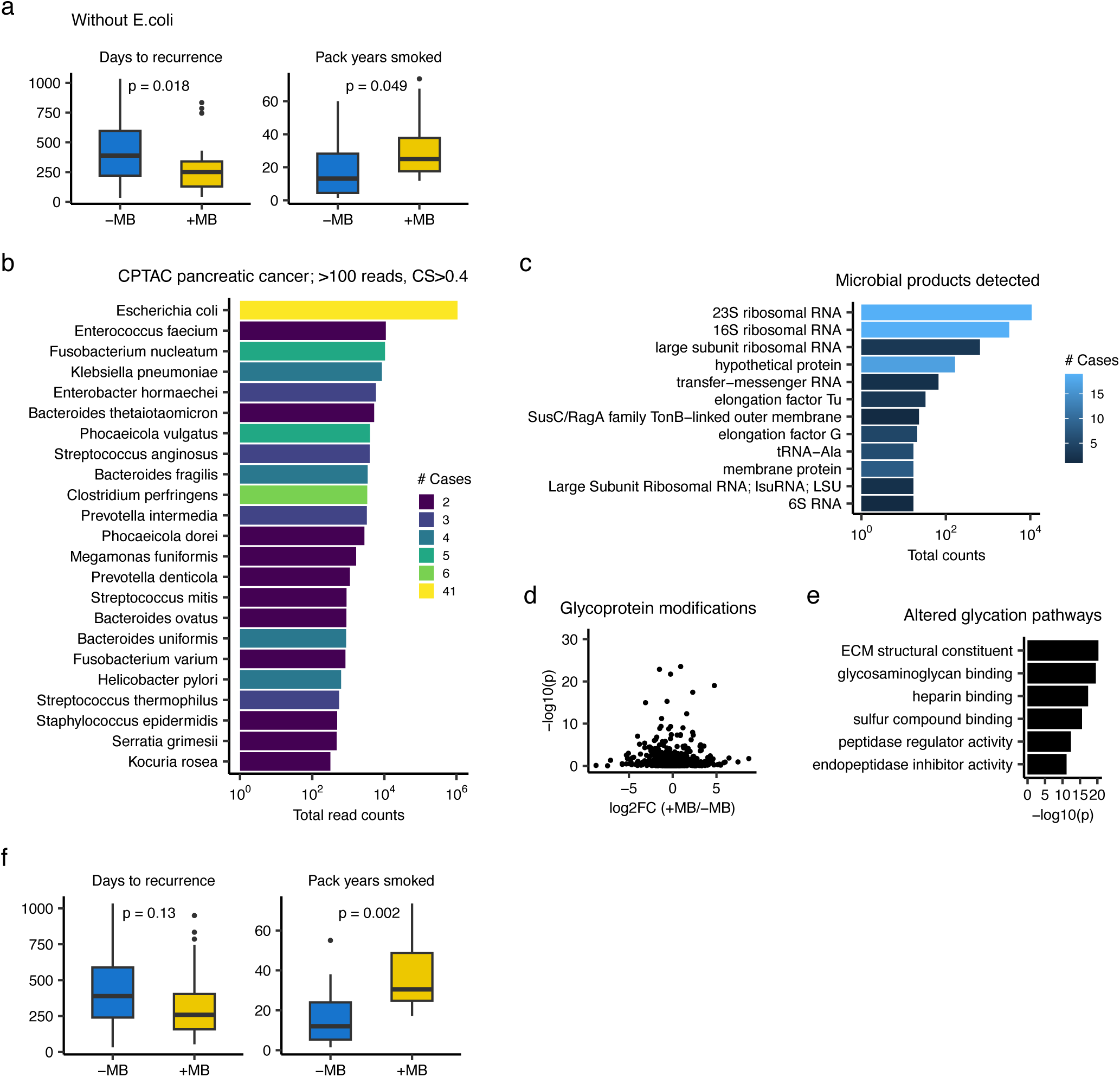
Detecting microbes in pancreatic cancer using a stricter PRISM score. **(a)** Boxplots of the days to tumor recurrence (left) and pack years smoked (right) of pancreatic cancer patients whose tumors have (+MB) or do not have (-MB) detectable microbes. Data were assessed with *E. coli* reads removed. Boxplots show median (line), 25^th^ and 75^th^ percentiles (box) and 1.5xIQR (whiskers). Points represent outliers. P-values are from Wilcoxon testing. (**b)** Bar plot indicating the total read counts of species identified in 159 cases of pancreatic cancer in CPTAC using a more stringent PRISM score (CS) of >0.4. Bars are colored by the number of cases in which the species was detected. **(c)** Bar plot of the most common microbial products detected in CPTAC pancreatic cancer. **(d)** Volcano plot of genes with glycoprotein modification. The x-axis is the log_2_-fold change of average gene modification level in tumors with detectable microbiome (+MB; n=91) vs. those without (-MB; n=68). The Y-axis indicates the Wilcoxon p-value. **(e)** Bar plot indicating gene ontology (GO) pathways of the genes with significantly different glycoprotein modifications (p<1e-10) in tumors with vs. without detectable microbiome. (**f**) Boxplots of the days to tumor recurrence (left) and pack years smoked (right) of pancreatic cancer patients whose tumors have (+MB) vs. do not have (-MB) detectable microbiome. Boxplots show median (line), 25^th^ and 75^th^ percentiles (box) and 1.5xIQR (whiskers). Points represent outliers. P-values are from Wilcoxon testing.

## Supplementary Tables

**Table S1.** CLID datasets

**Table S2.** Human reads missed by Kraken and STAR

**Table S3.** PRISM training and testing datasets

**Table S4.** PRISM validation datasets

**Table S5.** CPTAC and TCGA microbes

**Table S6.** 16S vs RNA-seq data profiled by PRISM

**Table S7.** Ontologies of altered glycoproteins in PDAC with vs. without detected microbes

**Table S8.** Compositions of the CLID-C and WGS datasets

